# A strong start for sustained success: inclusivity through a national group mentorship program for first-year graduate students

**DOI:** 10.64898/2026.03.12.710679

**Authors:** Sergio R. Labra, Valerie A. Tornini, Maria Pia Rodriguez Salazar, Daniela M. Cossio, Rebekah A. Gelpí, Bryan E. Rubio Perez, Yanitza M. Rodríguez, Gerardo Leana Sandoval, Kimberly Hernandez, Olivia V. Goldman, Robert W. Fernandez

## Abstract

In the United States, STEM graduate programs and workforce do not represent the demographics of the population. Obstacles, including a lack of transparency, community, and accessible information in navigating academia, disproportionately affect students from underserved backgrounds. Peer mentoring networks can address these disparities. Here, we describe Cientifico Latino, Inc.’s Graduate Student Engagement and Community (CL-GSEC) program, a nationwide, group-based peer mentorship program that has served first-year graduate students across the U.S., especially those from underserved backgrounds. Surveys indicate CL-GSEC positively impacts the first-year graduate experience. We highlight key program features, challenges, and insights, such as financial strains faced by first-year graduate students. We offer suggestions for how faculty and departments can better support students during this critical early stage of graduate training. We hope that reporting on CL-GSEC’s program structure, evaluations, and findings will guide educational leaders in expanding programming for junior graduate students.

## Introduction

The first year of graduate school is a pivotal moment for students pursuing Science, Technology, Engineering, and Math (STEM) careers. Although successfully admitted into competitive STEM graduate programs, many students experience feeling out of place and imposter syndrome; challenges that disproportionately affect those from underserved backgrounds (Clance and Imes 1978; Hall and Burns 2009; Dancy and Jean-Marie 2014; Fraenza 2016; Markle et al. 2022; Anonymous 2025). Research indicates that graduate students from underserved backgrounds who leave their graduate program do so in the first two years (Sowell, Allum, and Okahana 2015), and women are more likely to drop out during their first year of graduate school without adequate support systems (Foy 2018). With only 57% of graduate students finishing their PhD in 10 years (Sowell, Zhang, and Redd 2008), early support and intervention is critical for shortening graduate training timelines and improving graduation rates.

The challenges first-year students face often extend beyond psychological barriers. Many earn stipends below living wages (McClain 2025; Gaynor and Rautsaw 2025) and typically relocate across state or international borders away from their established support networks. Simultaneously, they must navigate the “hidden curriculum”, or the expectations and norms of academic culture (B. Smith 2013; Kentli 2009; Hariharan 2019; Pensky et al. 2021); all while developing their research directions, applying for fellowships, and searching for lab placements. These and other vulnerabilities caused by inconsistent student resources, funding, and mentorship are amplified for students from underserved communities, such as those with first-generation or lower socioeconomic family backgrounds (Cadena et al., 2023). Since the 2023 United States Supreme Court ruling prohibiting race-based affirmative action, the inclusion of scientists from underserved backgrounds in professional fields and higher education is expected to further decline (Bleemer 2020; Santoro 2023). Recent federal interventions targeting diversity, equity, and inclusion (DEI) programming and principles threaten to widen this gap further (White House 2025; Hill et al. 2025).

In this manuscript, we discuss an intervention program for supporting first-year graduate students in STEM fields. Between 2020 and 2025, Científico Latino, Inc.’s Graduate Student Engagement and Community (CL-GSEC) initiative has served 296 first-year graduate student scholars. CL-GSEC integrated a small-group mentorship structure along program-wide resources, webinars, and social events, to help support students through the financial, cultural, and professional transition of starting graduate school. Supplementing the community, professional networks and resources available at their home institutions, our goal was to set up first year students from underserved communities with a strong start to their graduate school, aimed towards increased retention and satisfaction in their STEM graduate school journeys and careers.

### Persistent underrepresentation and attrition of diverse STEM graduate students

Diversity of perspectives and expertise yields greater potential for scientific progress (Malcom 1996; National Academies of Sciences, Engineering, and Medicine 2023; Hofstra et al. 2020; Graves et al. 2022). However, ethnic, and racial, gender, ability, and socioeconomic diversity of the STEM workforce, as previously defined by the United States (U.S.) National Institutes of Health (National Institutes of Health, 2019), does not currently represent the United States population. Comprising 19.1% of the U.S. population, the number of Hispanic or Latino doctorate recipients in Science and Engineering lags at 9% (NSF Survey of Earned Doctorates, 2022). These numbers are similarly behind for Black or African Americans constituting 6% and American Indian or Alaska Natives receiving 0.5% of these doctoral degrees, despite them representing 12% and 1.1% of the population, respectively (NSF Survey of Earned Doctorates, 2022). Throughout the STEM academic career track, this disparity continues to increase, particularly at the faculty level (Wood et al. 2020). Contributing factors are more deeply felt by scientists from underserved backgrounds during pivotal career transitions, such as embarking on graduate studies, engaging in postdoctoral training, and undertaking the hiring process for leadership (Valantine, Lund, and Gammie 2016; Hoover 2023).

Graduate training is a pillar contributing to the development of STEM workforces. Globally, graduate student satisfaction with their programs is dramatically dropping (Nature Research 2022; Woolston 2022) and a mental health crisis in graduate education is well documented (Evans et al. 2018; Bekkouche, Schmid, and Carliner 2022). Challenges related to graduate school have a more acute effect on students from underserved backgrounds (Posselt 2021), defined here as individuals whose socioeconomic, ethnic, gender backgrounds are not equitably represented in academia and higher education. Despite targeted interventions, the retention and completion rates of STEM graduate students from underserved backgrounds lags behind those of students from privileged backgrounds (Gibbs 2014; Council of Graduate Schools 2024). Students from underserved backgrounds, such as Black and first-generation students, are reported to have higher attrition rates in graduate programs (Kniffin 2007; Nguyen et al. 2023). In one study, about half of graduate students from underserved backgrounds who left their graduate programs did so within the first two years (Sowell, Allum, and Okahana 2015).

### First year of graduate school is an intervention point for increasing support and belonging for students from underserved backgrounds

The challenges of transitioning into graduate school can be compounded for students from underserved backgrounds (Posselt 2021; Bekkouche, Schmid, and Carliner 2022). At the start of graduate school, all students must navigate a series of foundational milestones, including going through coursework, finding suitable research mentors, and preparing for PhD candidacy examinations. These initial steps set the stage for the rest of a student’s graduate training. However, ethnicity, gender, socioeconomic status, and other factors associated with underserved backgrounds often impose additional challenges in building mentoring relationships that foster awareness and confident understanding of the implicit expectations, norms, and knowledge of academia that are rarely taught directly. Lack of familiarity with this so-called “hidden curriculum” may leave students from underserved backgrounds feeling underprepared, isolated, or inadequate (B. Smith 2013; Kentli 2009; Hariharan 2019; Pensky et al. 2021). Furthermore, it may hinder their ability to integrate into academic culture and their departmental, laboratory, or peer cohort communities (B. Smith 2013; Park et al. 2022; Estrada et al. 2011; Dickens 2021; Bekkouche, Schmid, and Carliner 2022), or even contribute to students feeling that their personal values conflict with those of the U.S. academic system (McGee 2020).

First-year graduate students often relocate away from familial and community support networks, which students from historically marginalized backgrounds may rely on more heavily (Hinton, Termini, et al. 2020; Stachl and Baranger 2020). Furthermore, gender, racial, and citizenship status discrimination persists in academic environments and negatively affects graduate student mental health (Bernard et al. 2017; Posselt 2021; Woolston 2022; Nature Research 2022; Lee 2021). When entering graduate programs with limited ethnic or racial representation, new students may find it particularly difficult to build peer and mentor support groups that can recognize and empathize with their experiences. At the same time, faculty and graduate students from underserved backgrounds are often subject to the “minority tax”; disproportionally burdened with mentorship obligations and DEI-related institutional work (Rodríguez, Campbell, and Pololi 2015; Jimenez et al. 2019; Gewin 2020; Trejo 2020). This additional strain not only affects them but can reduce the time, energy, and resources available for them to provide effective mentorship. Ultimately, these factors shape the students’ well-being, sense of belonging, and received mentorship quality, all of which are important predictors of retention in STEM academia (Santiago and Einarson 1998; Inzlicht and Good 2006; Thomas, Willis, and Davis 2007; Griffith 2010; Fisher et al. 2019; Loyola and Grebing 2022).

The first year in graduate school is a key stage where mentorship can increase success that compounds over a student’s entire tenure in graduate school. Factors that support early graduate student success such as financial support, career opportunities and better mentorship could be an effective intervention (Nature Research 2022; Woolston 2022). Fostering social support and peer mentorship programs, with administrative support, are relatively easier to implement and can readily improve graduate student well-being (Lorenzetti et al. 2019; Charles, Karnaze, and Leslie 2022; Barreira and Bolotnyy 2024; Hopkins et al. 2024).

### CL-GSEC and peer mentorship as critical support for first-year graduate students

Mentorship is essential at each stage of academia, but culturally and racially relevant mentoring, particularly as it relates to the hidden curriculum of STEM academia, is critical to success, especially for graduate students from underserved backgrounds at the start of their graduate careers (Hinton, Vue, et al. 2020; McGee 2020; Pfund et al. 2022). Peer mentoring networks are an effective response to the lack of institutional support and existing disparities that may disproportionately affect students from underserved backgrounds in academic training (Xu 2008; Lorenzetti et al. 2019). Often, the hidden curriculum of academia is passed on informally by peer mentors, especially in community-oriented settings that aim to make STEM more accessible (Cadena et al. 2023; Bayin, McKinley, and LaFave 2023; Hinton, Vue, et al. 2020; Hopkins et al. 2024). A growing body of research suggests that students from underserved backgrounds, compared to their peers, possess forms of community cultural and social wealth, including peer support (Cadena et al. 2023; McGee 2020; Bourdieu 1986; Espino 2014; Stanton et al. 2022). These strengths can be supported and enhanced to make STEM more meaningfully inclusive.

To help first-year graduate student retention and well-being, we implemented a national peer mentorship program to support first-year graduate students in STEM. Here, we describe the Científico Latino, Inc.’s Graduate Student Engagement and Community (CL-GSEC) program, a U.S. nationwide group-based peer mentorship network aimed at supporting first-year graduate students of all backgrounds, especially those from underserved backgrounds. The goal of GSEC is to offer support beyond individual institutions and local networks, allowing incoming graduate students in search of community and mentorship to build a support system and a network of scientists from underserved backgrounds that may be lacking in their institutions. Since 2020, CL-GSEC has supported 296 first-year students from 121 (primarily U.S.-based) institutions.

CL-GSEC programming is built around the diversity of experiences and intersectional challenges among individuals within and across underserved backgrounds (Thomas, Willis, and Davis 2007; Curry et al. 2012). CL-GSEC programming addresses challenges first-year students face, such as applying to fellowships, navigating financial challenges, adjusting to academic expectations and norms, and, for students in rotation-based programs, choosing a research advisor.

CL-GSEC offers both a small group mentorship structure, which we call “mentorship pods” (Murrell and Onosu 2023), and a larger community network. Each mentorship pod comprises multiple mentors and mentee first-year participants (or “scholars”) as a way to bring about diverse perspectives in a small, safe, and inclusive setting. The larger network of the CL-GSEC community serves as a further resource. Importantly, the advancement of digital communication tools, including social network platforms, video-conferencing tools, and organizational messaging tools (e.g., Slack) provides an accessible and equalizing force for students from underserved backgrounds, especially those seeking community and information. The social acceptance and adaptation of asynchronous communication has enabled both more expansive networks of mentors and scholars from underserved backgrounds, and generally yielded a broader reach of networks beyond an institution or region (McReynolds et al. 2020). A decentralized, virtual, national mentorship program such as CL-GSEC enables participants to discuss more freely issues they may face as students from underserved backgrounds. Additionally, cross-institutionally connected students can exchange advice on how to navigate common challenges, without institutional oversight that may otherwise silence or exclude members (Hinton, Vue, et al. 2020; Jones-London 2020).

In this report, we share the design, implementation, outcomes, and insights gained from CL-GSEC. We focus on program data from the 2021-2022 and 2023-2024 academic cohorts, comprising 158 first-year students from 90 unique institutions, and 111 mentors from 66 unique institutions. Our data suggests that students from various underserved backgrounds may benefit the most from small group peer mentorship in their first year of graduate school. We hope others may adapt these observations and strategies for the benefit of their own communities.

## Results

### Implementation of a national first-year peer mentorship program and flexible team structure for scientist volunteers

The authors developed CL-GSEC to support students navigating the hidden curriculum of their first year in graduate school (Figure 1A). Components of the hidden curriculum addressed included financial costs and planning, applying for fellowships, time and project management, establishing mentorship networks, as well as navigating rotations, immigration-based challenges and social and cultural professional norms. CL-GSEC targets three key routes of support: Mentorship (e.g., through small mentorship groups), Community (through local and virtual social/networking opportunities), and Engagement (sustaining active participation in the program through features such as workshops) (Figure 1B). CL-GSEC aims to offer first-year graduate students support, community and mentorship beyond their home graduate program. This allows them to build and access connections and network wider than their local network. CL-GSEC’s potential for social impact in supporting first-year graduate students and fostering an inter-institution community are outlined in Figure 1C.

**Figure 1.**
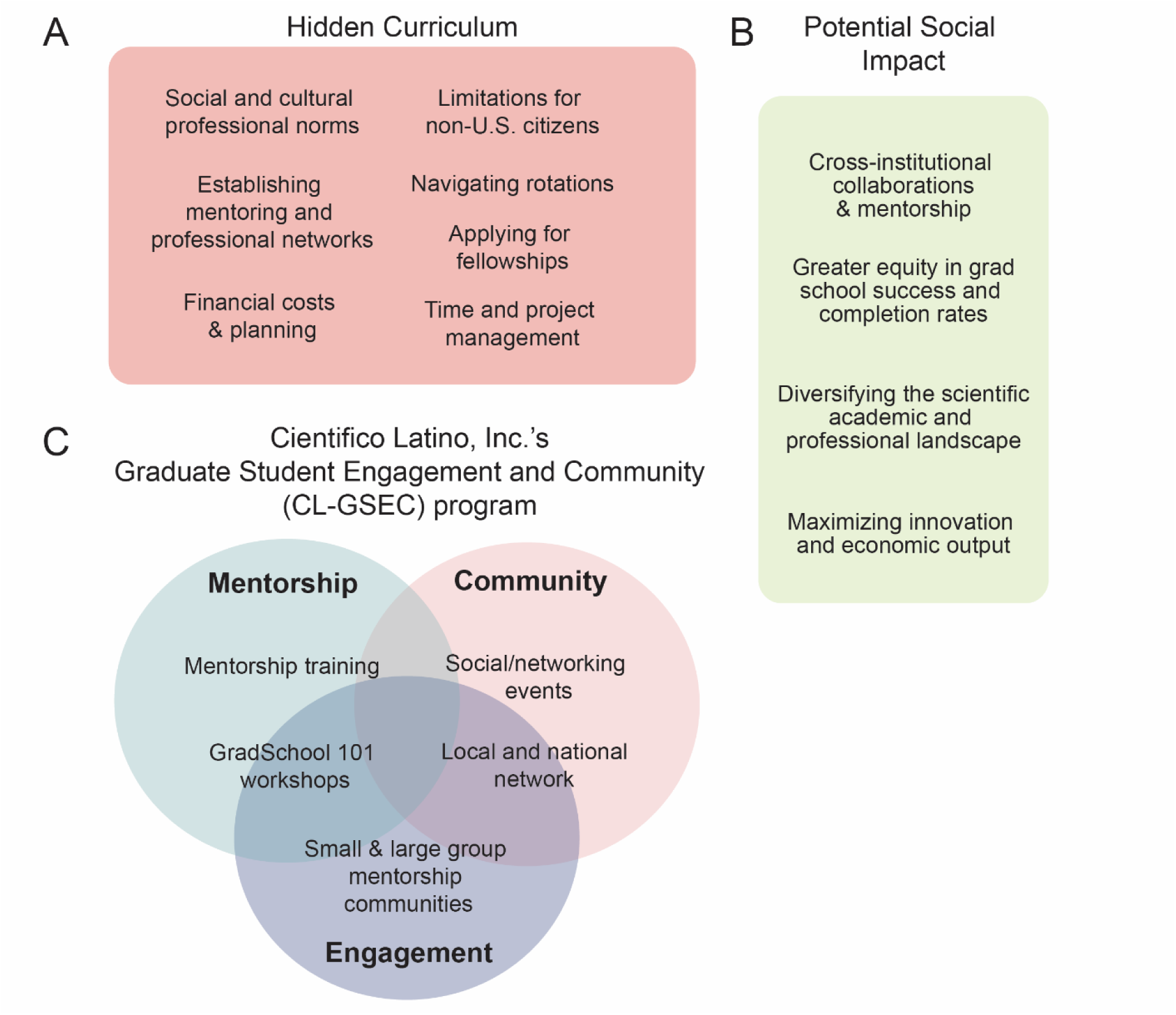
CL-GSEC program aims to create social impact through addressing the “hidden curriculum” and cultivating mentorship, community and engagement. **(A)** Cientifico Latino, Inc.’s Graduate Student Engagement and Community (CL-GSEC) aims to address components of the “hidden curriculum” for first-year graduate students. **(B)** CL-GSEC supports first-year graduate students through Mentorship, Community, and Engagement. Specific goals are provided for each. **(C)** Potential social impact of CL-GSEC, defined as significant or positive changes that address social injustices or challenges. This applies throughout students’ graduate school tenure and as they progress in their scientific training and careers.

CL-GSEC began as a semester-long pilot in spring 2021. It was then expanded to last the entirety of the academic year, and accepted cohorts for 2021-2022, 2023-2024, and 2024-2025. CL-GSEC took a hiatus in the 2022-2023 academic year to evaluate program structure, analyze data and improve the program based on scholar, mentor and team member feedback.

Accomplishing the goals of CL-GSEC required establishing a robust and flexible team structure. This allowed for the contribution and leadership of 20 team members and volunteers who were otherwise employed as full-time graduate and postdoctoral researchers. CL-GSEC’s executive leadership consists of two or three co-directors sharing the responsibility for preparing, executing, and evaluating the program. The executive team oversees and coordinates the operations of five other teams: Recruitment, Community Engagement, GradSchool 101 Workshops, Marketing, and Data teams (Figure 2). Their primary and inter-related functions are described in Supplementary File 1. This organization allows for CL-GSEC leadership and volunteers to effectively distribute program responsibilities and focus on their primary tasks. It also allows the flexibility for teammates to rely on each other during periods of time when, as graduate and postdoctoral researchers, they need to focus additional attention on their own career’s pressing milestones.

**Figure 2.**
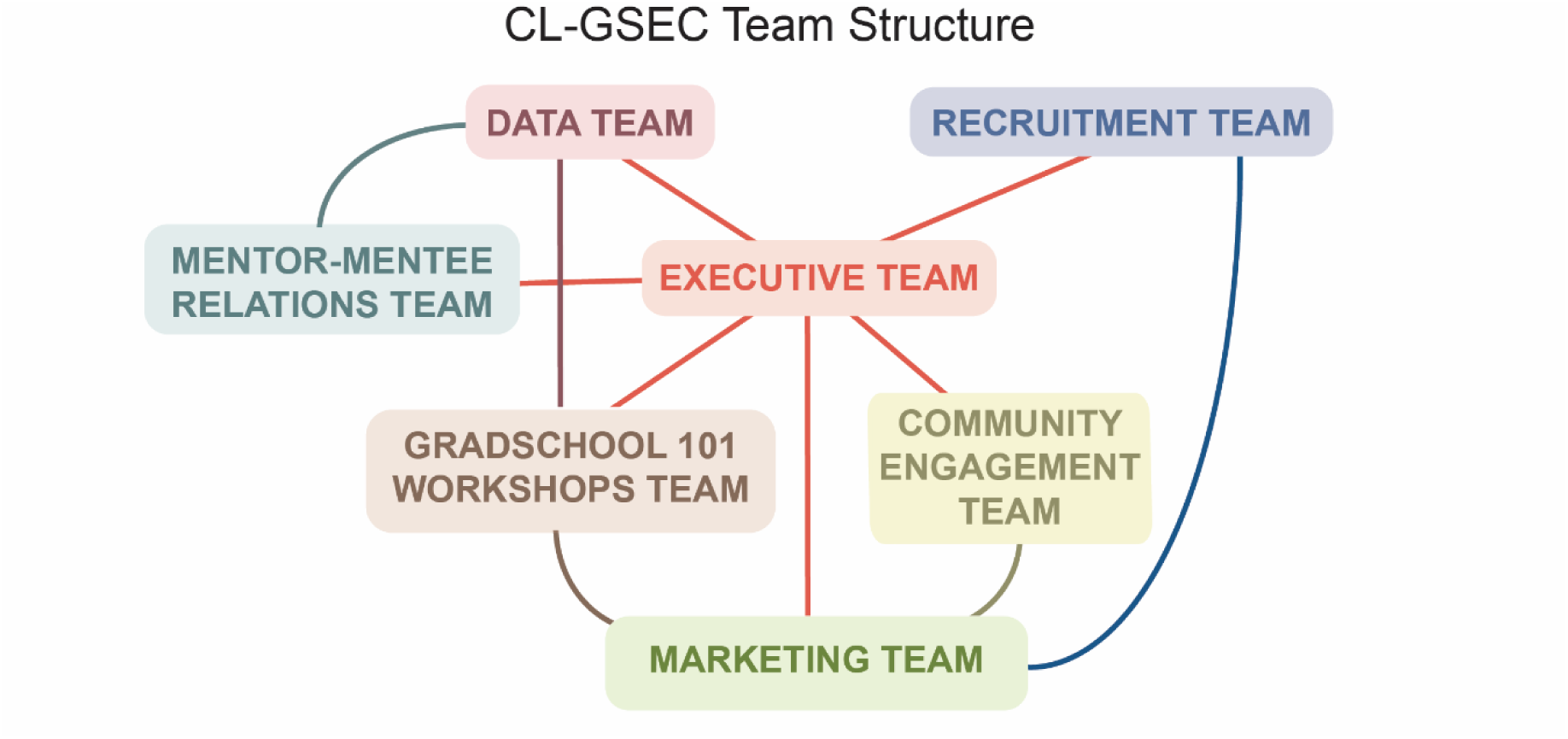
CL-GSEC program interconnected team structure allows for effective responsibility distribution and flexibility for researcher volunteers. CL-GSEC team structure and interconnectedness. Descriptions, functions, and size/management of each team are presented in Supplementary File 1.

### CL-GSEC comprised of students and mentors from diverse ethnic, racial, and scientific backgrounds

CL-GSEC scholars are first-year graduate students from institutions across the U.S., especially students from underserved backgrounds, as self-defined by race, gender, sexual orientation, ethnicity, socioeconomic status, and/or first-generation college student status. Before 2023, CL-GSEC scholars were recruited from an alumni pool of matriculated students who had previously participated in Científico Latino Inc.’s mentorship program for STEM graduate student applicants (CL-GSMI, detailed in Cadena et al. 2023). In 2023, CL-GSEC opened for public applications and accepted individuals other than CL-GSMI alumni. Mentors and scholars were recruited through the Científico Latino, Inc.’s website, newsletter, and social media (e.g., Twitter/X and Instagram).CL-GSEC recruitment aimed to attract applicants from diverse backgrounds. Emails were also sent out to graduate programs, deans, and DEI offices, including at minority-serving institutions, in attempts to reach eligible program applicants (incoming graduate students). Throughout CL-GSEC, many students who applied self-identified as “underrepresented”; no applicants were excluded from selection and thus all applicants were admitted to the program. In total, scholars across the three cycles of CL-GSEC represented 96 institutions, and mentors represented 71 institutions.

To understand our population of scholars, and to provide them with the mentorship they needed, we collected scholars’ preferences for attributes in their mentors as part of our application process. When signing up for CL-GSEC, scholars were asked to rank the relative importance of having specific features in common with their CL-GSEC mentors; options included field of study, preferred background identities, or geographical proximity (indicating the likelihood of meeting in person). Scholars ranked the field of study as their highest priority feature in their GSEC mentors, followed by mentor identity and geographic proximity (Figure 3A, Zenodo Supplemental Data: https://doi.org/10.5281/zenodo.18992301). We created scholar-mentor groups (“mentorship pods”) to best reflect this ranking.

**Figure 3.**
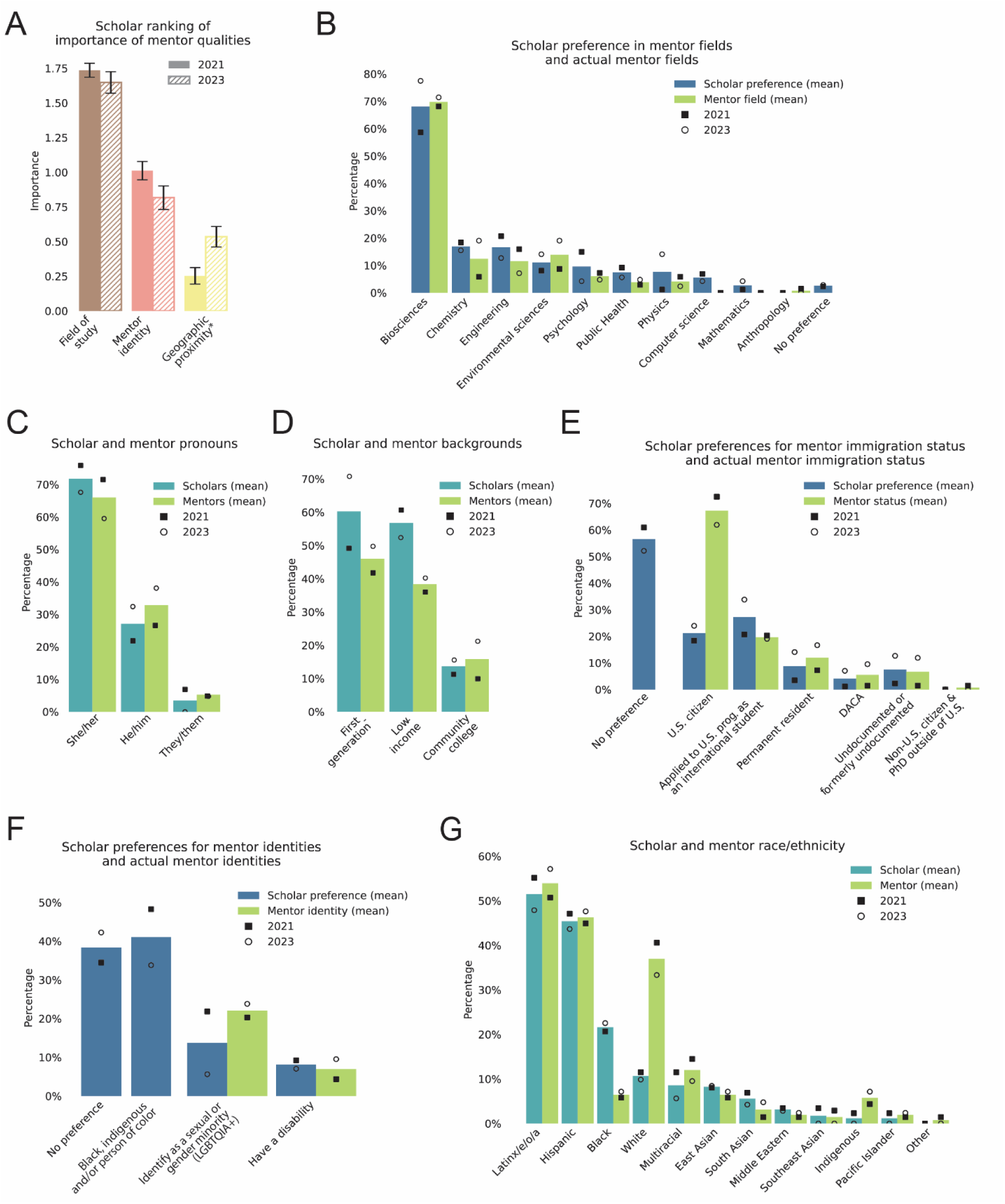
Scholar and mentor demographic information and features used in matching strategy for mentorship pods in CL-GSEC. **(A)** Scholars ranked the relative importance of mentor qualities. Each option was ranked as “Most important”, “Important”, or “Least important”, here represented as 2, 1, and 0 respectively for comparison. Data from 2021 (solid bars) or 2023 (hatched bars) cohorts. Error bars indicate standard error of mean (S.E.M.) **(B-G)** Proportion of scholars or mentors that selected features listed below barplot. Features indicate scholar preference in mentor identity (blue), scholar self-identification (teal), and mentor self-identification (green). Data from 2021 (square) or 2023 (circle) cohorts. Bars indicate mean proportions from both cohorts. **(B)** Scholar preference for mentor field of study compared to mentor field of study (respondents could select multiple fields). **(C)** Scholar and mentor pronouns (respondents select from a set or write-in their responses). **(D)** Scholar and mentor socioeconomic and educational backgrounds. **(E)** Scholar preference for mentor immigration status compared to mentor immigration status (respondents could select multiple fields). **(F)** Scholar preference for mentor identity features compared to mentor identity features. Both scholars and mentors could select multiple fields. “Black, indigenous and/or person of color” was not an available option for mentors. **(G)** Scholar and mentor race/ethnicity (respondents could select multiple fields). For exact questions, response options, survey responses and scripts to generate all figures, see Zenodo Supplemental Data. 2021 cohort, n = 87; 2023 cohort, n = 71. *“Geographic proximity” was changed to “Same or nearby institution” in 2023.

We recruited prospective mentors that were either graduate students (in at least their second year of studies and/or advanced to PhD candidacy) or postdoctoral researchers. Although CL-GSEC’s mentor recruitment did not target any specific fields of study, selected mentors happened to represent a similar distribution of scientific fields of study per scholars’ STEM field preferences (Figure 3B). As indicated by mentor preferences, most of our scholars were enrolled in graduate programs in Biosciences (58.654.0% in 2021 and 77.56.1% in 2023), Chemistry (18.416.1% in 2021 and 15.5% in 2023), or Engineering (20.719.5% in 2021 and 12.7% in 2023) (Figure 3D). Accordingly, CL-GSEC mentors were recruited in matching fields of study (68.166.7% in biosciences in 2021 and 71.4% in 2023; 5.8% in chemistry in 2021 and 19.1% in 2023; 15.917.4% in engineering in 2021 and 7.1% in 2023) (Figure 3D). This compatible recruitment structure enabled us to form appropriate small mentorship groups (two mentors, four scholars).

For assessing CL-GSEC scholar preference for mentor identity, we took into account features including gender identity, family and educational background, immigration status, race and ethnicity, as well as others (Figure 3C-G). CL-GSEC scholars and mentors were recruited in similar proportions in several of these categories, such as self-identified pronouns and backgrounds. The majority of scholars used “she/her” pronouns (75.9% in 2021 and 67.6% in 2023) indicating our program primarily served students who identified as women (Figure 3C). Moreover, scholars and mentors came from low-income backgrounds (Scholars: 60.9% in 2021, 52.6.2% in 2023; Mentors: 36.2% in 2021, 40.5% in 2023), were first-generation college students (Scholars: 49.4% in 2021, 71.1% in 2023; Mentors: 42.0% in 2021, 50.0% in 2023), and some had previously attended community colleges (Scholars: 11.5% in 2021, 15.8% in 2023; Mentors: 10.1% in 2021, 21.4% in 2023) (Figure 3D). We also took into account immigration status, as this can have significant effects on first-year students’ personal experiences in adjusting to their graduate programs and academic factors such as eligibility for fellowships, particularly for international and undocumented students (Lee 2021; National Science Foundation, 2025; National Institutes of Health, 2025). We were able to recruit mentors across different immigration statuses including U.S. citizens, international students, DACA, among others, matching roughly to scholar preferences (Figure 3E).

The CL-GSEC program attracted a diversity of historically underrepresented identities in STEM. Among CL-GSEC scholars, a large proportion requested mentors that identified as persons of color (48.3% in 2021, 33.8% in 2023). The majority of CL-GSEC scholars self-identified as “Latinx/e/o/a” (55.2% in 2021 and 47.9% in 2023), “Hispanic” (47.1% in 2021 and 43.7% in 2023), and “Black” (20.7% in 2021 and 22.5% in 2023) (Figure 3G).CL-GSEC mentors self-identified in similar proportions, although with less mentors identifying as “Black” and more “White” (“Latinx/e/o/a”: 50.7% in 2021 and 57.4% in 2023; “Hispanic”: 44.9% in 2021, 47.6% in 2023; “Black”). Because of CL-GSEC’s group mentorship structure, scholars could have multiple mentors with diverse identities. Mentors whose identity-informed perspectives were in higher demand could easily support multiple scholars.

### Mentorship pods for allows for increased flexibility, diverse perspectives for mentors

A key element of CL-GSEC is a small group mentorship structure (or “mentorship pods”). Mentorship pods consist of 2 senior graduate student mentors and 4 first-year graduate students per group. This differs from the 1:1 mentorship structure commonly employed by other programs (for example, Científico Latino, Inc.’s GSMI (Cadena et al., 2023), <u>Muse Mentorship</u>, <u>National Research Mentoring Network</u>, <u>Project Short</u> and <u>SACNAS MAS program</u>) in allowing for flexibility and division of labor between mentors within a mentorship pod.

There were 11 small groups in the 2021 pilot, and approximately 20 small groups in each subsequent iteration of CL-GSEC. Initial monitoring of mentorship pods took place through feedback forms. These were intended to provide limited supervision for, as well as identify and resolve issues, for scholars and mentors across small groups. In the pilot year, feedback forms were requested on a monthly basis. Subsequently, this was revised to once every 2 months based on participant evaluation. After each check-in survey, we identified groups with low satisfaction or participation notes and connected with them to identify the sources of problems and possible solutions. To account for scholar or mentor non-participation, groups were rearranged as needed, for example combining two smaller, actively participating groups together.

We assigned the membership of each mentorship pod to maximize balance of expertise and ability to find common meeting times. To aid in pod membership assignment, we took into account scholars’ needs and preferences, as well as scholar and mentor and mentors fields of study, identities, and geographic region (Figure 3). This group-based mentorship structure allows exposure to diverse perspectives at a critical step of academic training. Moreover, group-based mentorship also creates an immediate built-in community that includes both mentors and peers.

To support both mentors and scholars in the program, mentorship resources and training were provided (Supplementary File 2). Scholar resources covered topics such as effective mentee practices, the importance of flexibility and consideration, self-advocacy, active engagement in meetings, aligning expectations with mentors, and preparing meeting agendas. Mentor resources included advice on interpersonal skills and communication, sponsorship, communicating expectations, and ways to engage and connect among mentorship groups. Suggested discussion topics relevant to a typical U.S. academic year cycle were provided to mentors to help facilitate monthly meetings (Supplementary File 3).

### Community-based curriculum and activities to address first-year graduate student challenges

Graduate school is one of the most stressful stages of scientific training (Levecque et al. 2017; Evans et al. 2018; Welsh 2022). The COVID-19 pandemic further exacerbated isolation, especially for first-year graduate students that were geographically separated from their families, friends, and communities (Ogilvie et al. 2020). Finding and building community and a sense of belonging are crucial to wellbeing and success in higher education and graduate school (Walton et al. 2023; Shepard and Perry 2022; Johnson 2022; Tay 2021). The group-based mentorship program is supplemented by local and cohort-wide activities. These include workshops and virtual and in-person socials, hosted by CL-GSEC team members or mentors, to facilitate the exchange of perspectives, building of community, and sharing of mentorship resources. Three to four large-group virtual socials are planned throughout the program. CL-GSEC also hosted monthly virtual coffee hours, for informal discussions outside the small mentorship groups. We also included a small number of coffee hours that were Spanish-inclusive, driven by mentor and scholar requests for opportunities to connect in a common experience and language other than English. In-person meetups were implemented 2023-2024 CL-GSEC cycle, often taking place in metropolitan regions where mentors and scholars are concentrated or during larger conferences (including those hosted by Society for Neuroscience, American Society for Cell Biology, and the Society for the Advancement of Chicanos/Hispanics and Native Americans in Science).

Lastly, the CL-GSEC program provides a webinar series to discuss challenges of the first-year of graduate school, called “GradSchool 101” (Figure 4). We designed these webinars to focus on critical topics and challenges in the hidden curriculum for first-year graduate students (Figure 4A). The webinars consist of panel discussion with volunteer mentors and team members (Figure 4B). Topics included “First-year Fellowships,” “Choosing a Lab,” “Time Management,” and “Setting Expectations.” High interest in these topics has led to them being offered each year of CL-GSEC, along with newer panel offerings on “Imposter Syndrome,” “Tax Preparation,” and “Public Speaking.” Webinars distributed throughout the academic year and timed for relevance. For each webinar, we surveyed scholars before and afterwards to learn how the workshops could be improved and more tailored to their needs.

**Figure 4.**
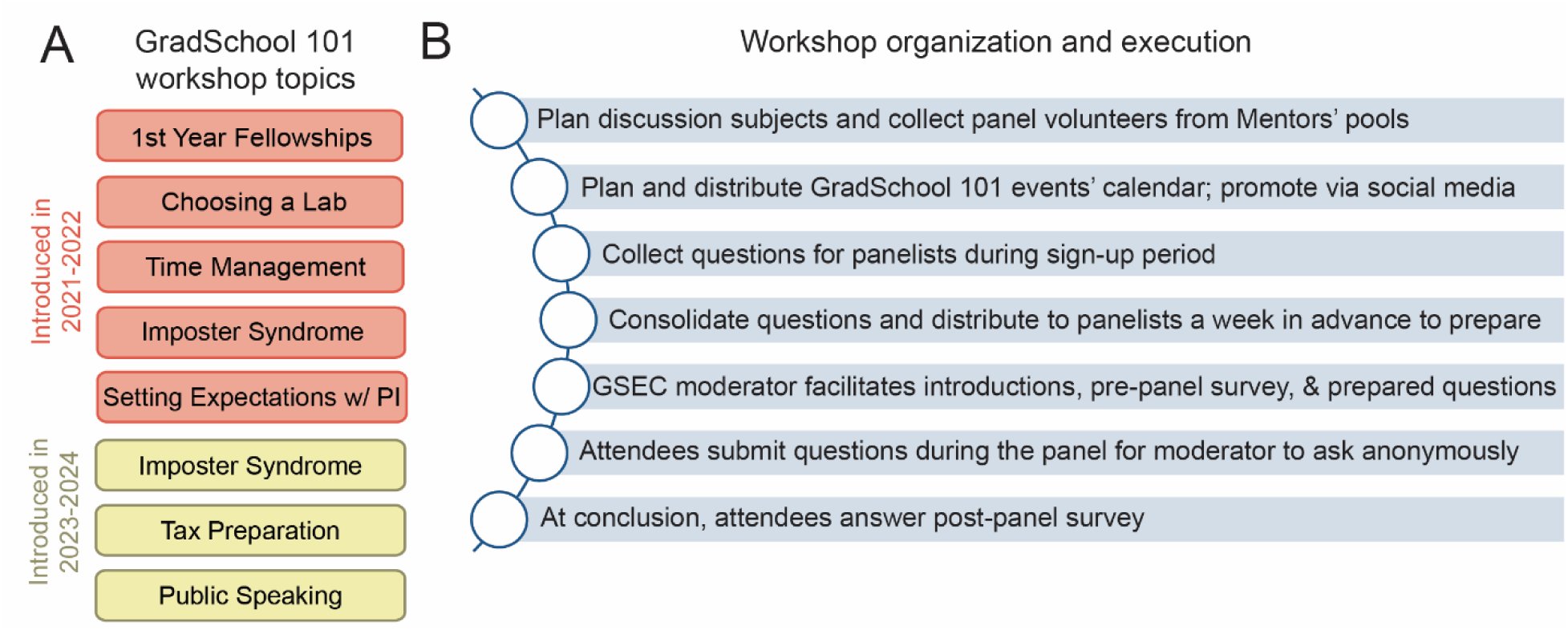
CL-GSEC GradSchool 101 workshops to address first year graduate student needs. **(A)** CL-GSEC GradSchool 101 Workshop topics. Topics offered in the 2021-2022 academic year included: First year fellowships, choosing a lab, time management, and setting expectations with Principal Investigator (PI). Imposter syndrome, tax preparation, and public speaking were added in the 2023-2024 academic year. **(B)** CL-GSEC GradSchool 101 organization, execution, and internal evaluation protocol.

Overall, CL-GSEC community-wide activities supplemented the mentorship and discussion from their mentorship pods, and provided a wider network and sets of expertise on which students could turn to for support.

### Positive impact of CL-GSEC program determined by participant and mentor evaluation

At the end of the program, CL-GSEC scholars and mentors were asked to answer an evaluation questionnaire regarding their experience with the program and their small mentorship groups. On average, about half of the total cohort filled out the final evaluation surveys (46% in 2021; 55% in 2023). Of survey respondents, over 90% of scholars responded that CL-GSEC helped them navigate their first year of graduate school (90% in 2021 and 92.3% in 2023; Figure 5A). On a scale from 1 to 5 (minimum to maximum satisfaction), the majority of scholars ranked the program with a 4 or higher (4.35±0.12 in 2021, and 4.20±0.14 in 2023; Figure 5B). The large majority of CL-GSEC scholars expressed an interest in becoming future program mentors (70% in 2021; 84.6% in 2023), demonstrating their conviction in the program’s impact on future scholars (Figure 5C).

**Figure 5.**
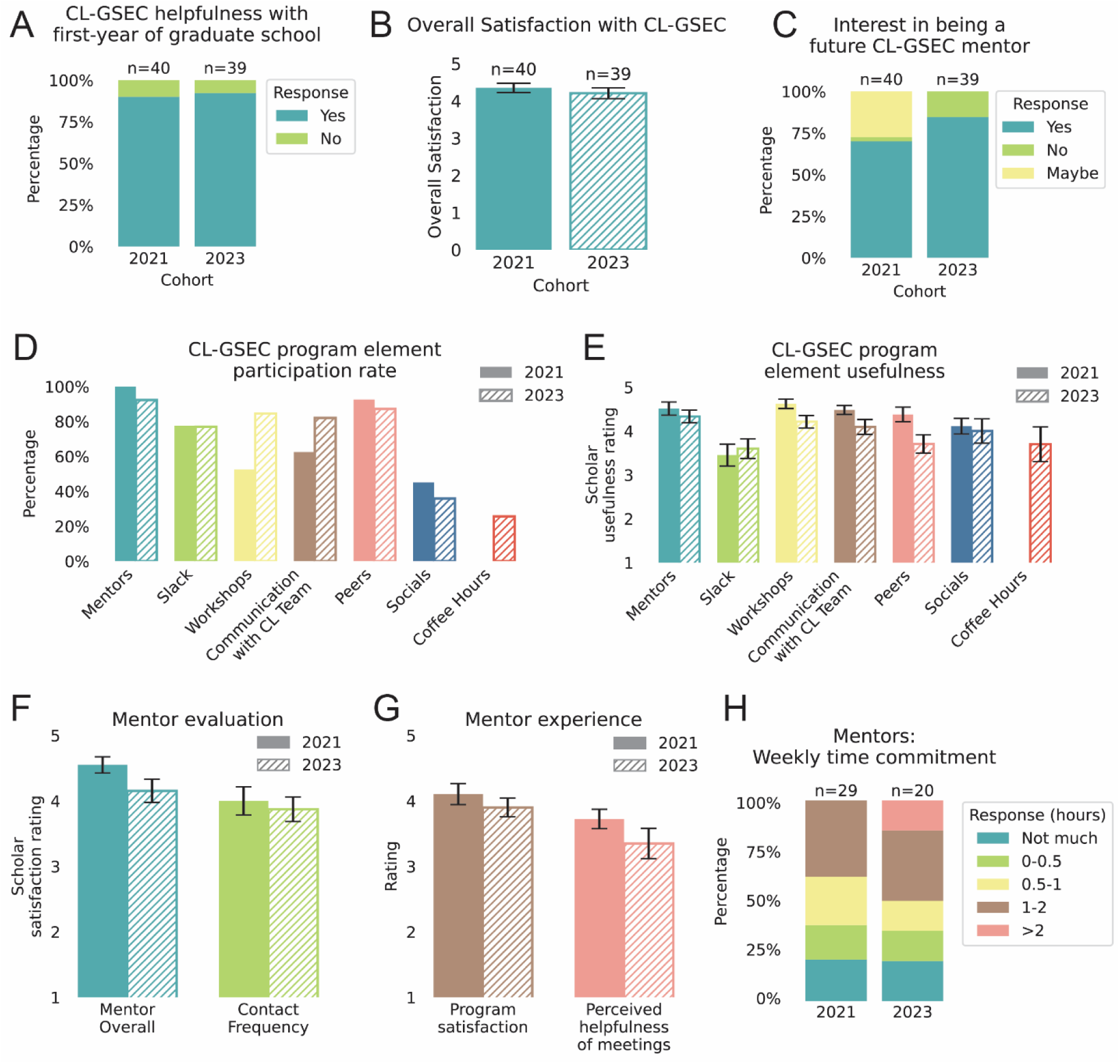
CL-GSEC scholar and mentor program evaluation and outcome. **(A-E)** GSEC scholar responses to end of program survey. Figures illustrate data collected from 40 respondents from the 2021 cohort (out of 87) and 39 respondents from the 2023 cohort (out of 71). For exact questions, response options, survey responses and scripts to generate all figures, see Zenodo Supplemental Data. **(A)** Proportion of respondents for each cohort that answered “Did being part of GSEC help you navigate your first year of graduate school in any way?” with “Yes” (teal) or “No” (green). **(B)** Overall scholar satisfaction with the CL-GSEC program. Respondents on a scale from 1 to 5 (minimum to maximum satisfaction). Bars represent mean and error bars indicate S.E.M. **(C)** Scholar-indicated interest in becoming a CL-GSEC mentor (after passing qualifying exams). Respondents answered “Yes” (teal), “No” (green) or “Maybe” (yellow). For ease of analysis and mentor recruitment in subsequent years, the “Maybe” option was removed for the 2023 cohort. **(D)** Scholar participation of various elements of the CL-GSEC program including mentors, slack workspace, virtual workshops, communication with the CL-GSEC team, relationships with their peers, program socials, and coffee hours. Bars represent mean and S.E.M. (error bars). Coffee hours were first introduced with the 2023 cohort. **(E)** Bar plots showing self-reported participation among scholars for each of the program elements in D. The 2023 cohort was asked to rate usefulness on a scale from 1 to 5 (minimum to maximum usefulness), while the 2021 cohort was offered categorical responses (“Very useful”, “Somewhat useful” or “Could use improvement”, from which they could select multiple), which were then adjusted to a 1 to 5 scale for comparison across cohorts. **(F)** Scholar satisfaction of their assigned mentors and frequency of contact with them, ranked on a scale from 1 to 5 (minimum to maximum satisfaction). **(G-H)** GSEC mentor responses to end of program survey. Figures illustrate data collected from 29 respondents (out of 69 total) from the 2021 cohort and 20 respondents (out of 42 total) from the 2023 cohort. **(G)** Overall mentor satisfaction with the CL-GSEC program, and respondent answer to the question “How helpful do you think your monthly mentor meetings were to your mentees?”. Respondents answered on a scale from 1 to 5 (minimum to maximum satisfaction or helpfulness). Bars represent mean and error bars are standard error of mean (S.E.M.). **(H)** Proportion of categorized responses to the open-ended question “How much time do you feel this program required from you on a weekly basis?”. Open-ended responses were assigned to the following categories: “Not much” (teal), indicating unspecified or negligible time commitment, or in hours “0-0.5”, “0.5-1”, “1-2”, “>2” (indicated in green, yellow, brown, and pink, respectively). If a range was given, the higher value was used to categorize the response.

In the CL-GSEC closing survey at the end of the academic year, we asked scholars which program features they participated in and their perceived usefulness (Figures 5D and E). The mentors were among the most helpful agents of the program, with an average rating of (in 2021, 100% participation and a 4.51±0.15 out of 5 rating; in 2023, 92.3% participation and 4.33±0.14). Peer scholars (other first-year graduate students in their small mentorship groups) followed as a close second (in 2021, 92.5% participation and a 4.37±0.17; in 2023, 87.2% participation and 3.71±0.20). The Slack channel had relatively high participation (77.5% in 2021 and 76.9% in 2023), although lower rated usefulness (3.45±0.25 in 2021; 3.60±0.22 in 2023). The GradSchool101 workshops were ranked highly useful (in 2021, 52.3% participation and a 4.62±0.11; in 2023, 84.6% participation and 4.21±0.14), in addition to communications with the CL team (in 2021, 62.5% participation and a 4.48±0.10; in 2023, 82.1% participation and 4.09±0.17). Although less than half of scholars participated in the quarterly virtual socials (45% in 2021 and 35.9% in 2023), they were ranked useful among those that attended (4.11±0.18 in 2021; 4.00±0.28 in 2023).

To understand with more granularity scholars’ experiences in the mentorship pods, we asked scholars about their satisfaction and communication with their mentors. Overall, scholars showed a high level of satisfaction with their mentors (4.55±0.12 out of 5 in 2021; 4.15±0.18 in 2023) as well as with their frequency of contact (4.00±0.22 out of 5 in 2021; 3.87±0.18 in 2023) (Figure 5F).

We also surveyed mentors to understand their experience with CL-GSEC. Mentors felt overall satisfied in the program (4.10±0.16 out of 5 in 2021; 3.90±0.14 in 2023) (Figure 5G). They also perceived their small group meetings as helpful to their scholars (3.72±0.15 in 2021; 3.35±0.23 in 2023) (Figure 5G). At least half of mentors in each cohort reported their weekly time commitment as less than an hour per week (62.1% in 2021; 50.0% in 2023) (Figure 5H).

### Positive program experiences and points for improvement from CL-GSEC scholars and mentors

To improve the CL-GSEC program and better understand the needs of the community, we wanted to qualitatively understand their subjective experiences. In a free-response question, we asked CL-GSEC scholars and mentors to identify their most positive or helpful experience in the program, as well as identify negative experiences (Figure 6). Free-response questions were categorized responses to illustrate and evaluate the groups’ experiences (original responses are in Zenodo Supplementary Data).

**Figure 6.**
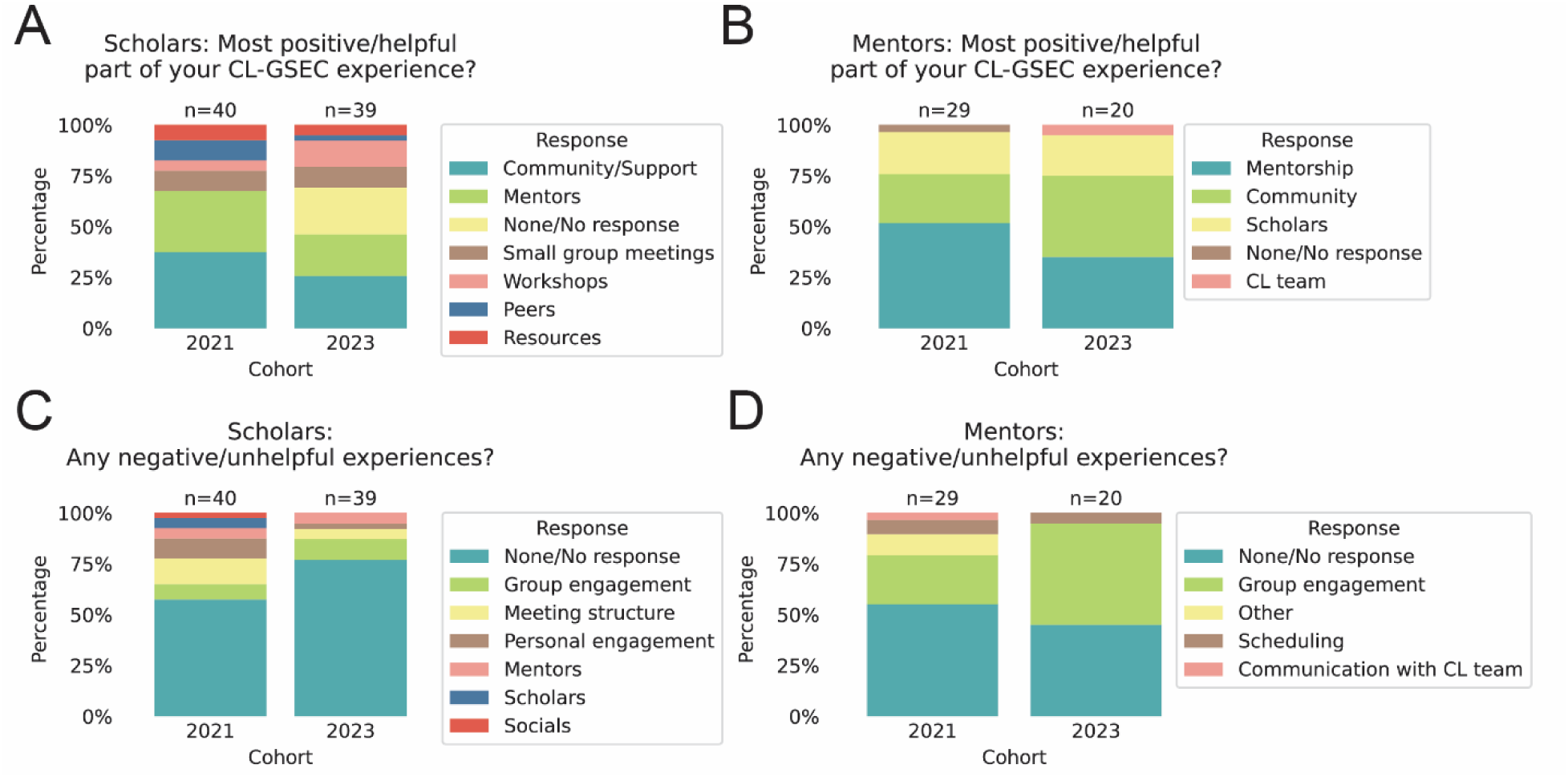
Qualitative responses and feedback from CL-GSEC scholars and mentors. Scholars and mentors were prompted for positive (A-B) and negative (C-D) experiences in CL-GSEC to derive a qualitative understanding of participant experiences. Number of responses indicated above bar. **(A-B)** Categorized responses to an open-ended question prompting scholars (A) and mentors (B) to provide positive program experiences. “If none, type N/A” was added to 2023 scholar questions for A and C, but was not included in 2021 questions or mentor questions. We have included exact questions in Zenodo Supplemental Data. **(C-D)** Categorized responses to an open-ended question prompting scholars (C) and mentors (D) to provide negative program experiences.

When asked about their most positive experiences, both scholars and mentors identified a sense of community and support (Scholars: 37.5% in 2021, 25.6% in 2023; Mentors: 24.1% in 2021, 40% in 2023), suggesting that this was a valuable supplement to their support network at their home institutions (Figure 6A-B). Other positive experiences reported by scholars included relationships with mentors and peer mentors, small group meetings, resources, and GradSchool 101 Workshops. Mentors, in addition, frequently reported a positive experience in being able to help graduate students more junior to them, categorized here as “Mentorship” (51.7% in 2021; 35% in 2023) (Figure 6B).

Representative positive experiences from scholars included:

> “The most positive part of my experience was having the opportunity to discuss the hardships and accomplishments of my first year of graduate school with a diverse group of scholars. As a Latina in STEM, it is often hard for me to find people who can understand and sympathize with my experiences so having a community of POC scholars allowed me to feel safe to express myself.”
>
> “I appreciated hearing from other mentees and also from mentors about different grad school experiences. Also found [it] helpful having someone outside of my bubble that I could reach out to.”
>
> “Feeling like I wasn’t alone in this journey”
>
> “[The] most positive part was getting together with the other mentees and mentors and [to] chat every month. Those interactions were the core of the program and I feel like they validated my experience as a grad student. Other mentees usually had similar experiences and we could talk our way through them, and mentors would often have useful feedback.”
>
> “I got the chance to share time and learn from my wonderful mentors and other mentees, get inspired by them, be vulnerable with them and learn that I am not alone in this experience.”

When mentors were asked “What was the most positive part of your experience in the CL-GSEC program” and “Overall, how was your experience in the CL-GSEC program?”, some answers included:

> “Getting to see the mentees encourage each other and seeing them actually talk to each other and reach out to us for help, meaning they trusted us”
>
> “I enjoyed being paired with a partner to help the burden of scheduling, or leading conversation. I really liked my mentees and that they all showed up for the group, and each other.”
>
> “The most positive part of my experience was being able to connect with mentees, provide support, and build community with them as they navigated their first year of graduate school.”
>
> “Being able to help individuals early in their graduate career by telling them about what I wish I had done differently and encouraging them to advocate for themself. I could see that being with peer mentors gave them some confidence when making big decisions.

We present additional de-identified feedback quotes from both CL-GSEC scholars and mentors in Supplementary File 4.

Each year, we asked mentors and scholars for feedback on improving the program, which was implemented whenever possible. For example, in response to mentors asking for more guidance and structure in the 2021 closing surveys, in 2023 we disseminated detailed mentorship training and guidance (Supplementary File 2) and suggested monthly topics (Supplementary File 3). Participant feedback has been critical for the development and refining of CL-GSEC’s effectiveness.

When asked “What do you wish the GSEC program had offered that you would have benefitted from?”, CL-GSEC scholars suggested: (1) additional workshops topics (such as on being a Person of Color in academia, time management, self-care); (2) smaller CL-GSEC gatherings at home/nearby/regional institutions; (3) more information and guidance about applying for fellowships (particularly for personal statements) and identification of fellowships, including for non-U.S. citizens; and (4) accountability or online writing groups.

When scholars were asked, “What additional resources or guidance would have been helpful for you, in regards to your relationship with your CL-GSEC mentors?”, scholars proposed: (5) structure to meetings, such as an outline of what would be discussed that month, and mentors bringing up questions (to avoid instances where scholars not really knowing what to ask); (6) more check-ins from CL-GSEC team to see if groups were meeting; (7) a short bio of mentors at the start of the program so they could more easily find common ground; and (8) setting expectations of how often they will meet at the beginning of the year.

Independently, CL-GSEC mentors suggested: (1) improving program structure to clarify scholar expectations and encourage their engagement; (2) for socials to occur more frequently and to be more structured (e.g., specific activities or topics); (3) more guidance for mentors, including resources and feedback; and (4) more diverse workshops topics. Lack of group engagement was a particular source of negative experience for mentors (24.1% in 2021; 50.0% in 2023). Concurrently, some scholars expressed that they wished they had engaged more with CL-GSEC programming (10.0% in 2021; 2.6% in 2023). Future iterations of the program addressed this by a mentor and mentee handbook outlining expectations of participants of this program; an orientation for mentors and mentees going over expectations and a program timeline for the semester; incorporating scientific training workshops (such as how to use computational pipeline tools, how to make scientific figures, how to read scientific papers, and preparing for non-academic careers); structured in-person and virtual socials at different universities and timezones; a shared Slack channel and cross-community social for all Científico Latino, Inc. programs to encourage community building and collaboration; and a scholarship fund for first-year graduate students to help them with educational expenses in their first year of graduate school.

When prompted for both positive and negative experiences, many more scholars and mentors provided positive responses than negative responses (Figure 6C-D). We note slight changes in question phrasing across years can impact results. For instance, “If none, type N/A” was added to 2023 scholar questions (Figures 6A and 6C), but was not included in mentor questions or 2021 scholar questions (Zenodo Supplemental Data). In summary, assessments from scholars and mentors were notably positive, indicating CL-GSEC’s effectiveness in building a supportive community to support first-year graduate students during this critical transition period.

### Effects of financial barriers on overall first-year experience of CL-GSEC scholars

We have previously reported that financial hurdles are a large challenge for graduate school applicants from underserved communities when applying to graduate school [Cadena et al., 2023]. We sought to understand the extent to which these challenges persisted in students’ first year of graduate school.

We surveyed the CL-GSEC 2023 cohort and program alumni (first-year graduate students in 2020-2021 and 2021-2022 academic years) to determine whether they faced financial barriers when transitioning to graduate school. An excerpt from a student about the costs of relocation to transition to graduate school:

> “I was a commuter student during my undergraduate, so I depended on my parents a lot for financial stability. While my parents supported me in chasing my dreams and moving out, they were not able to assist me financially to move to a new state for graduate school given the tight finances at home. Therefore, I depended a lot on savings and credit cards to afford my move. This includes unexpected costs such as purchasing all new furniture, apartment applications, housing supplies, first-month rent, security deposit, etc. These costs were a heavy burden for me given that I didn’t receive my first paycheck until the middle of September when by that time I already paid 2 months’ worth of rent. This experience introduced a lot of stress to me because I am still paying the repercussions of credit card debt.”

About a third of scholars from both groups reported no unexpected expenses (32.5% in 2023; 31.3% among alumni). The majority CL-GSEC scholars mentioned they experienced unexpected expenses towards laptops, access membership societies (e.g., Society for Neuroscience, SACNAS), textbooks, specialized software expenses (e.g., statistical analysis packages) in their first year of graduate school (among 40 scholars in the 2023 cohort: 30.0%, 25.0%, 12.5%, 2.5%, respectively; among 16 alumni scholars: 31.3%, 31.3%, 31.3%, 18.8%, 31.3%) (Figure 7A). Scholars estimated spending over $300 towards these expenses (2023 cohort: approximately $322; alumni: approximately $399) (Figure 7B).

**Figure 7.**
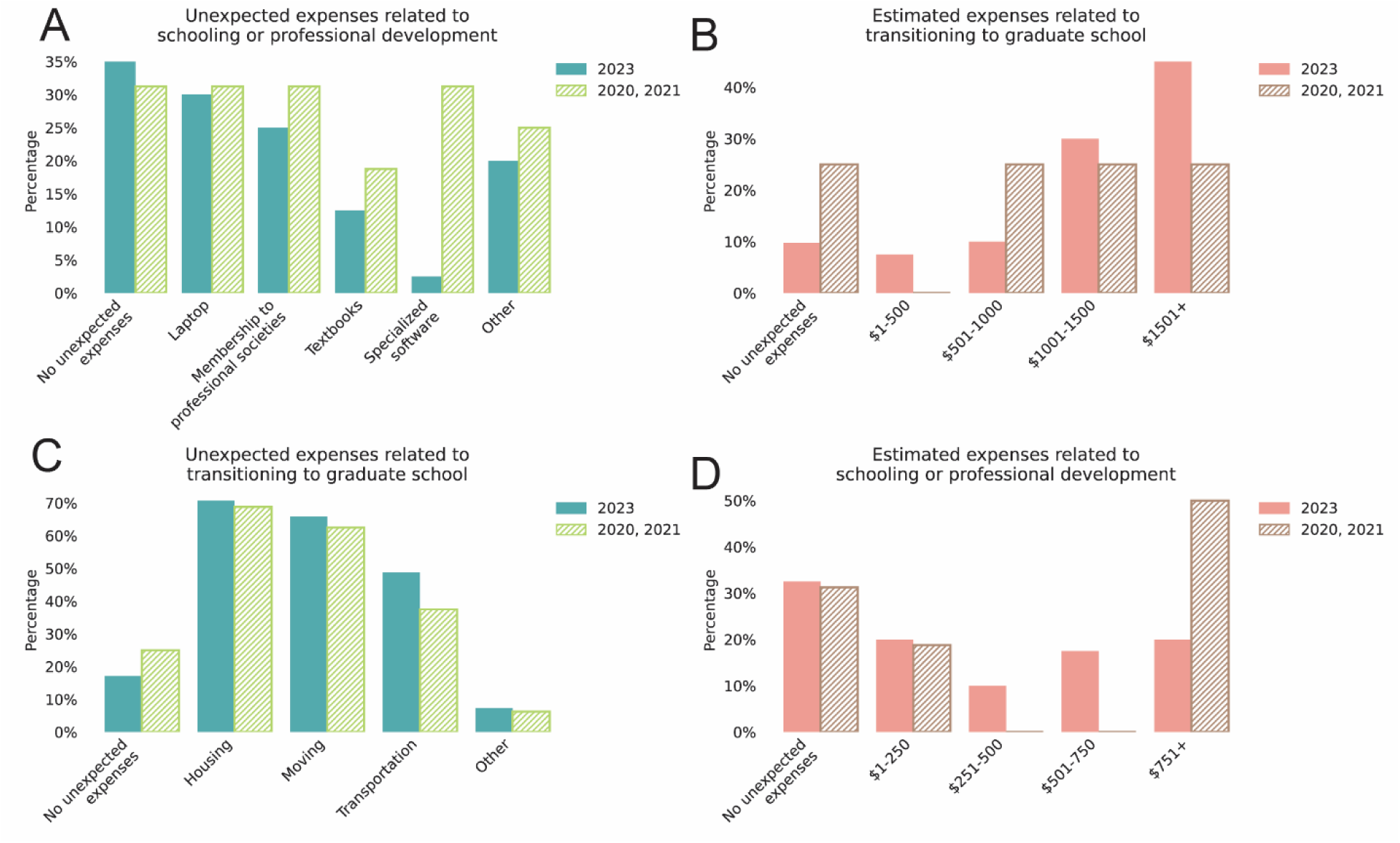
Financial expenses in the first year of graduate school. **(A-D)** Proportion of 2023 cohort scholars (solid bars) or program alumni from 2020 and 2021 cohorts (hatched bars) that self-reported expense type and amount related to starting graduate school. **(A-B)** Expense type (A) and estimated amount (B) related costs for schooling or professional development (2021 cohort, n=40; 2020 cohort, n=4; 2021 cohort, n=12). **(C-D)** Expense type (C) and estimated amount (D) related costs of transitioning to graduate school (2021 cohort, n=41; 2020 cohort, n=4; 2021 cohort, n=12).

Although it is common for first-year graduate students to move across state or even international borders for their graduate school education, the costs of relocation are rarely covered. Even more commonly than professional development, scholars often had costs related to relocation, such as housing, moving, and transportation (among 41 scholars in the 2023 cohort: 70.7%, 65.9%, 48.8%, respectively; among 16 alumni scholars: 68.8%, 62.5%, 37.5%) (Figure 7C). In 2023, 7.5% estimated spending between $1-500, 10% spending $501-1000, 30% spending $1001-1500, and 45% spending over $1501 (Figure 7D). Among alumni, 25% fell into each estimated spending category: $501–1,000, $1,001–1,500, and above $1,501. Taking into account that several scholars reported no unexpected expenses (9.8% in 2023, 25% among alumni), overall, scholars regularly spent at least $800 towards these expenses (2023 cohort: approximately $1100; alumni: approximately $875).

## Discussion

Graduate school and the process of scientific discovery are challenging, particularly when students do not feel supported (Nature Research 2022; Woolston 2022; Anonymous 2025). Underserved graduate students’ sense of belonging and retention can be influenced by factors such as relocation from family and community support, unfamiliar academic systems, inequities related to gender, race, and/or citizenship status, challenges around local peer and mentor networks, and financial strain (Santiago and Einarson 1998; Inzlicht and Good 2006; Loyola and Grebing 2022; Hinton, Termini, et al. 2020; Stachl and Baranger 2020; Bernard et al. 2017; Nature Research 2022; Lee 2021; Council of Graduate Schools 2024).

To address this, we developed CL-GSEC, a volunteer-run, small group-structured mentorship program for first-year graduate students from underserved backgrounds. Here, we analyzed program and evaluation data from CL-GSEC’s 2021-2022 and 2023-2024 academic cohorts, comprising 158 first-year students and 111 mentors from 124 unique institutions.

Scholars came from a range of scientific, socioeconomic, educational, race/ethnicity and immigration backgrounds. We created small groups, or “mentorship pods”, composed of 4 first-year graduate student scholars and 2 senior graduate students. This enabled scholars to receive diversity of perspectives from mentors and peers, and spread responsibilities between mentors, particularly for those of more underrepresented backgrounds. Mentorship pods were curated based on scholar-reported preferences. Through evaluation surveys, scholars and mentors reported program satisfaction and ranked the effectiveness of individual program elements, particularly mentors and workshops. In qualitative analyses, we identified that the most valuable component of the program for both scholars and mentors were the CL-GSEC community, interpersonal and group relationships. Participants, particularly mentors, also shared areas for improvement, such as a desire for increased mentorship pod engagement. We also explored the unique finances of this transition period in junior scientists careers, which can lead to financial inequity, an additional challenge for first-year students from low-income backgrounds.

### CL-GSEC provides complementary resources and support to those offered at students’ home institution

Overall, CL-GSEC has (1) established a supportive network of scientists, especially of those from underserved backgrounds; (2) increased first-year graduate students’ understanding of graduate school expectations; (3) developed and disseminated strategies for first-years during a critical year of graduate school. Now in its sixth iteration for the 2025-2026 academic year, CL-GSEC has evolved to best serve the needs of its community of scholars.

We conclude that CL-GSEC, a year-long, mentorship and community engagement programming independent from students’ graduate program institutions, provides complementary and focused support for first-year students, especially relating to the hidden curriculum of academia. We highlight the program components such as: open-access resources and workshops, virtual small support groups that offer safe environments for students to seek advice, peers and mentors who may themselves experience and/or understand the challenges that are exacerbated for students from various underserved backgrounds, and consistent check-ins to ensure scholars are supported. These features may contribute to first-year graduate students feeling more sufficiently supported, increased satisfaction in their programs and, eventually, retention and success of STEM students from underserved backgrounds.

Complementing home institutions’ programming and support, CL-GSEC equips scholars with resources and guidance to navigate the hidden curriculum of graduate school, fostering a strong foundation for an empowered start and long-term success. CL-GSEC helps scholars overcome the challenges of their first-year transition by providing mentorship, building community, and creating cross-institutional engagement between scholars and mentors.

### Mentorship programs fill common gaps in the support and retention of students from historically marginalized backgrounds

Group peer mentoring is crucial at every stage, including principal investigators (Greco et al. 2022). CL-GSEC implements a specialized group mentorship structure for first-years to address students’ sense of belonging and provide guidance on challenges related to marginalized identities and backgrounds. Our approach differs from emphasis on “personal resilience”, which is disproportionately asked of students from underserved backgrounds (McGee 2020). This is particularly important for students from underserved backgrounds who may not have access to mentors of similar background at their home institutions (E. P. Smith and Davidson 1992; Austin 2002; Thomas, Willis, and Davis 2007). We note that mentorship programs alone cannot counteract structural barriers faced by students from historically marginalized backgrounds. Advisors and administrators to graduate students share responsibility to practice evidence-based, culturally responsive mentorship that supports students’ wellbeing and success.

For example, as with other historically marginalized groups, CL-GSEC aims to address the unique difficulties faced by international students and undocumented students. These include learning new cultural and academic norms, lack of access to fellowships, or facing extra teaching requirements if they are ineligible for public grants from principal investigators. These students may need to seek other jobs to make ends meet. These students may also struggle to find community among those unfamiliar with their unique challenges. In providing a space for mentorship and community engagement, CL-GSEC increases students’ chances of finding information and support that they may not have been able to identify at their home institutions.

Impactful mentorship relationships benefit both mentees and mentors. This is evidenced by CL-GSEC mentors also finding community within the program and fulfillment in guiding first-years through experiences similar to their own. Group mentorship structures lessens the burden on mentors, demonstrated by the relatively light time commitment required by CL-GSEC mentors. This is important in minimizing the “minority tax”, a phenomenon where mentors from underserved backgrounds are often called upon to perform additional labor in guiding junior students on issues related to demographics, race, ethnicity and/or gender, which may in turn may strain their own productivity and wellbeing (Rodríguez, Campbell, and Pololi 2015; Jimenez et al. 2019; Gewin 2020; Trejo 2020). Group peer mentorship expands access to mentors, spreads the burden of mentorship, and offers an efficient model of mentorship for STEM academia.

In a time when universities may not be able to appropriately support their students, especially those from underserved backgrounds, CL-GSEC and similar programs may provide graduate students access support, mentorship, and community. Such programs include, but are not limited to, the <u>Black in X</u> networks, including <u>Black in Neuro</u> (Murray et al. 2021), <u>Black in AI</u>, <u>Black in Cancer</u>, <u>Black in Genetics</u>, and <u>STEMNoire</u>; <u>LatinXinBME</u>; <u>DisabledInStem</u> Mentorship program (Paparella 2024); <u>500 Queer</u> <u>Scientists</u>; <u>500 Women Scientists</u>; American Indian Science and Engineering Society (<u>AISES</u>); Society for the Advancement of Chicanos/Hispanics and Native Americans in Science (<u>SACNAS</u>); <u>SPARK Society</u>; and the Biology Undergraduate and Master’s Mentorship Program (BUMMP) at University of California, San Diego (Ravishankar et al. 2024). By working with and learning from these and other initiatives, we hope to continue shaping CL-GSEC and learning how to best support students across different cultural and multi-identity backgrounds while positively recognizing and affirming their value (Dahlberg, Byars-Winston et al. 2019).

### Future improvements to CL-GSEC: program participation and tracking long-term impact

Among CL-GSEC survey respondents, scholars and mentors were overall satisfied with their participation and in open-ended responses, indicated positive experiences related with the community and their program interactions. In addition to more direct suggestions, such as workshop topics, participants indicated negative experiences. Some participants and mentors were disappointed at times in their groups’ lack of group engagement. At the same time, some scholars indicated a disappointment in not having engaged more with CL-GSEC programming. This indicates some scholars would have liked more engagement, while other scholars would have liked to engage more however felt it was beyond their bandwidth, likely due to adapting to high expectations and work demands of graduate school. We note that this represents a paradox among first-year graduate students who both need and have limited free time to accept help and support.

This is a persistent difficulty in non-mandatory and long-term support programs geared for graduate students. In CL-GSEC, participation drop-off was a common occurrence each year, particularly in the spring semester, which was addressed by re-consolidating scholars and mentors which participated into new mentorship pods. Scheduling difficulties for mentorship pods were also a persistent problem, which we addressed in 2021 by making sure mentorship pods were in similar timezones. Other possible solutions could be to provide an incentive structure for program engagement (e.g., gift cards or award recognition), or limiting participants to students and mentors with a known track history for engaging in similar programming (such as CL-GSMI).

Although we obtained metrics for scholars’ first-year experience, future evaluation of CL-GSEC scholars can address how participation translates to long-term and professional success. For the 2023 CL-GSEC cohort, all 39 closing survey respondents were still enrolled in their Master’s or PhD program by the end of their first-year, with one respondent expressing intent to transfer programs (Zenodo Supplemental Data). Future follow-ups and qualitative reflection from CL-GSEC scholars could enable us to gain a better understanding of the kinds of support that were most impactful. Further research could contribute to research on imposter syndrome, a key phenomenon that disproportionately affects students from underserved backgrounds, and elucidate interventions that specifically help first-year graduate students build confidence (Clance and Imes 1978; Hall and Burns 2009; Dancy and Jean-Marie 2014; Fraenza 2016; Markle et al. 2022). More research and parsing of first-year graduate students’ experience and intervention strategies would improve CL-GSEC as well as STEM graduate programs overall.

### First-year students experience acute financial strain

Our survey data indicated that first-year graduate students face high relocation costs and housing expenses. First-year graduate students are also generally responsible for education-related costs such as purchasing a new laptop, transportation to their college or research laboratory, textbooks for their courses, and specialized software to conduct their research (e.g. statistical software or programming platforms).

Without adequate support for incoming graduate students, students experience additional financial stress that can interfere with their academic potential throughout their PhD. Some graduate students may work second or third part-time jobs to make ends meet to lessen the financial opportunity costs of being in graduate school (Sims 2021). Additionally, graduate students may go to food banks to save on their food expenses (Graham 2024). These financial challenges are especially exacerbated for underrepresented students, with a report indicating that 49% of Black STEM PhD graduates in STEM fields complete their degrees in debt (Velez and Heuer, 2023).

### Suggestions for more equitable and accessible first-year graduate school retention and success

We present some recommendations for the faculty and departments which can be implemented to support graduate students in Figure 8. A low-cost and meaningful step towards providing support includes clarity and accessibility of information published on school or lab websites, such as funded research opportunities, lab expectations, mentorship dynamics, or a pathway to connect with program/lab alumni for career guidance. We also encourage departments to establish structured mentorship programs, implement STEM outreach awards, and provide professional development workshops on tackling imposter syndrome and navigating alternative careers to help improve the graduate student experience and career preparation. Support at the department and the university level includes providing subsidized housing, stipends for books and laptops, large relocation costs, and access to food banks. Universities can also host annual graduate student symposiums for students from underserved backgrounds to build community across different departments and establish cross-institutional partnerships with minority-serving institutions (MSIs) to aid with graduate student recruitment and representation. We note that some institutions, such as Rockefeller University (Rockefeller University) and University of Michigan (University of Michigan), address some of these issues through generous relocation costs, subsidized housing, stipends for academic expenses, and even reduced food costs.

**Figure 8.**
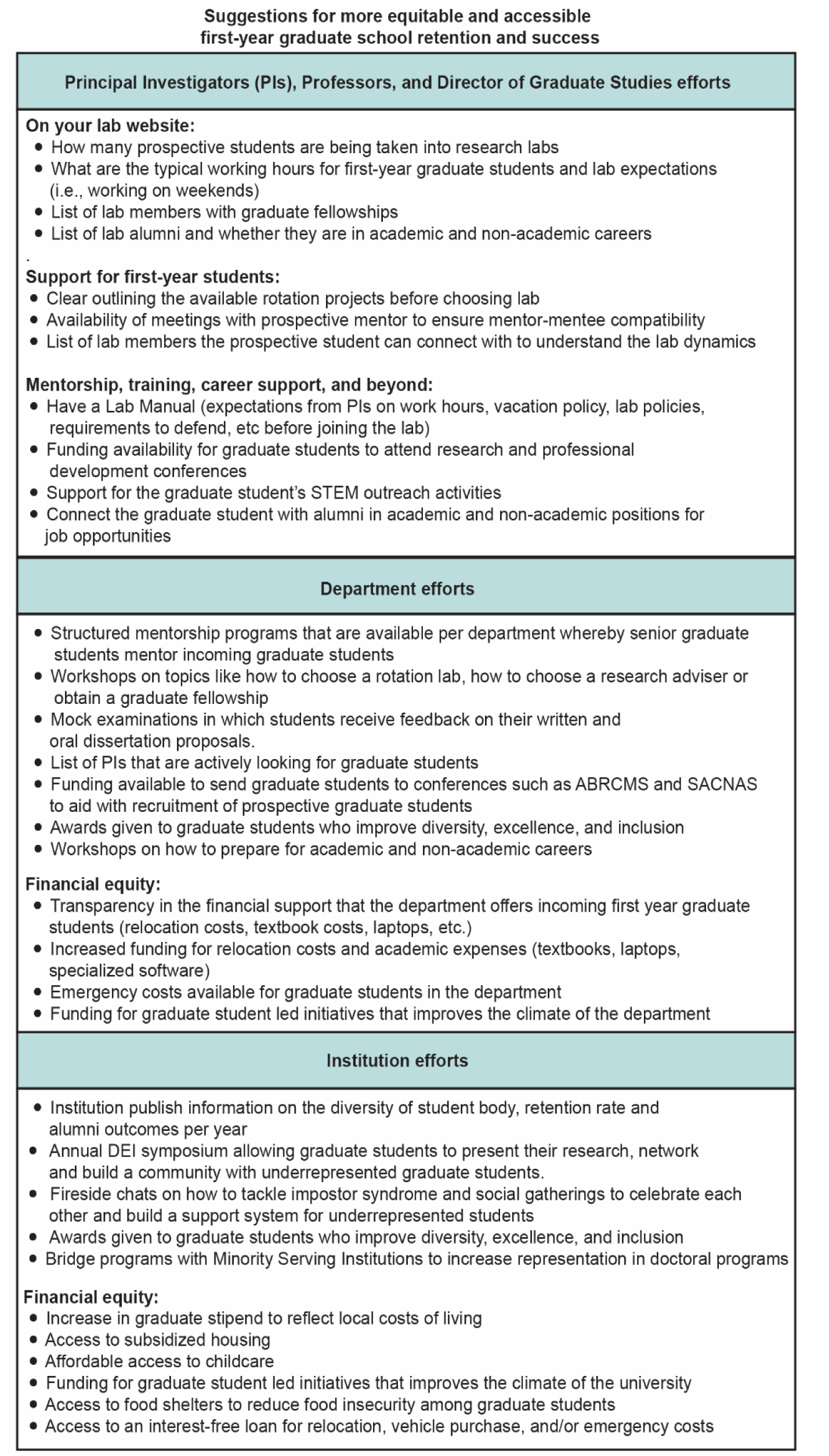
Suggestions for more equitable and accessible first-year graduate school retention and success.

### CL-GSEC offers a blueprint for an independent support network, free from institutional constraints

Our data, the personal experiences of the manuscript authors, and research referenced in this manuscript overwhelmingly suggest that current graduate school support systems do not adequately or equally support students across backgrounds. STEM higher education must actively create infrastructure and affect academic and departmental culture to enable the inclusion and success of graduate students from all backgrounds.

Recent events since the 2023 U.S. Supreme Court ruling prohibiting race-based affirmative action are poised to lead to the further exclusion of scientists from underserved backgrounds in professional fields and higher education (Santoro 2023). For example, diminishing of visible efforts to make STEM graduate student bodies reflect the general U.S. population could engender feelings of isolation or exclusion among students from underrepresented backgrounds, potentially diminishing their ability or enthusiasm for pursuing graduate studies and post-graduate academic or scientific careers.

Furthermore, State and federal government has made efforts defund, dismantle, and disable educational support systems, including funding mechanisms (e.g. Diversity Supplements) and impactful programs to make STEM graduate education accessible, such as Maximizing Access to Research Careers (MARC), Post-Baccalaureate Research Programs (PREP), Initiative for Maximizing Student Development (IMSD), and NIH’s Postbaccalaureate Program (Chronicle Staff 2024; Executive Order 14151 - “Ending Radical and Wasteful Government DEI Programs and Preferencing”; Office of Management and Budget Memo M-25-13). Finally, the end discretionary funding to Minority-Serving Institutions (MSI) grant programs will again limit critical research and institutional inroads to a more inclusive higher education system (Sauer 2025; U.S. Department of Education, 2025). At time of writing and publication, several of these orders still await legal scrutiny and legislation. All the while, the same economic, social, and educational disparities persist, further constraining the opportunities available to students from underserved communities further, rendering the pursuit of graduate studies daunting, less appealing, and more inaccessible.

In this manuscript and our description of the CL-GSEC program, we provide a blueprint for a community-led and -focused program that takes account of and responds directly to the needs of graduate students, especially those from underserved backgrounds, for individual success and for an inclusive scientific enterprise in the U.S. and beyond. Despite the rapidly changing landscape for this work in the United States, CL-GSEC and similar programs can operate on local or national scales, independently of university infrastructure which may be under institutional jurisdictions to state or federal mandates. We hope that the results and guidance provided can be implemented by leaders in science aiming to develop similar systems of support towards retention.

### Concluding thoughts

CL-GSEC offers resources, mentorship and community to foster a sense of belonging among first-year graduate students. We outline our program structure and outcomes for the benefit of those interested in carrying out similar work. We also offer suggestions to faculty and departments for improvements towards a mentorship and workforce training efforts that supports students from underserved backgrounds and would benefit all students, regardless of background. These efforts bring us closer to an equitable, inclusive, and accessible academia which in turn benefits all of society.

## Supplementary Files

**Supplementary File 1**. Descriptions of each CL-GSEC team, their general function, and their size/management.

**Supplementary File 2**. Training resources for CL-GSEC scholars and mentors.

**Supplementary File 3**. Suggested topics for monthly meeting discussions during a typical U.S. academic year.

**Supplementary File 4**. Additional testimonials from CL-GSEC scholars and mentors.

## Data and code availability

Survey questions and data from CL-GSEC scholars and mentors (anonymized, without identifying features), plots, analysis, and custom scripts used for figure generation are available at Zenodo (https://doi.org/10.5281/zenodo.18992301). Any additional information required to reanalyze the data reported in this paper is available from the corresponding authors upon request.

## Methods

The data collection and use of all surveys in this study were reviewed and determined by the Scripps Research Institute’s Institutional Review Board (IRB) to be exempt from formal committee review under 45 CFR 46.104(d)(2). CL-GSEC was composed of 45 scholars and 22 mentors in 2020-2021 (spring semester pilot program), 87 scholars and 69 mentors in 2021-2022, 71 scholars and 42 mentors in 2023-2024. We use data from the program’s closing survey, which was collected from 40 out of 87 scholars, and 29 out of 69 mentors in the 2021-2022 cohorts. Closing survey responses for the CL-GSEC program were collected from 39 out of 71 scholars, and 20 out of 42 mentors in the 2023-2024 program. For the research purposes of this manuscript, identifying features were removed, and all data was anonymized before analysis. For exact questions, response options, survey responses and scripts to generate all figures, see Zenodo Supplemental Data.

## Acknowledgments

We thank the CL-GSEC mentor volunteers who made and continue making this program possible and the CL-GSEC scholars for their feedback on this program. We thank Kimberly E. Leon, Valeria Montserrat Juarez, Lear Brace, Wendy Aquino Nunez, Anika Pruthi, Gabriel Lozano Betancourt, Julio Fierro, and Jose Luis Guerra for their contributions to CL-GSEC during their time on the team, and Dr. Laura Nicholson (Scripps Research Translational Institute, La Jolla, CA) for her support in acquiring IRB compliance.

CL-GSEC in the 2024-2025 academic year was supported by the Richard Lounsbery Foundation and a Biomedical Ad Hoc grant (1345547) from the Burroughs Wellcome Fund and, including the drafting and completion of this manuscript. S.R.L. was supported by the NIH TL1 program (TR002551), the Scripps Research Institute’s Kelly group, and the ARCS Foundation; R.A.G. by Canada Graduate Scholarship-Doctoral (CGS-D) from the Social Sciences and Humanities Research Council of Canada; M.P.R.S. by the Paul and Daisy Soros Fellowship and the Howard Hughes Medical Institute Gilliam Fellowship; D.M.C. by a NIH T32 (AG00096-40) and UCI Faculty Mentor Program Fellowship; B.E.R.P. by the University of Wisconsin-Madison Graduate Engineering Research Scholars (GERS) Program; K.H.P. by the Ford Foundation Pre-Doctoral Fellowship. O.V.G was supported by the Kavli Neural Systems Institute Graduate Fellowship, the National Science Foundation Graduate Research Fellowship Program and the Schmidt Science Fellowship; R.W.F. by a grant from the Simons Foundation (855199) and a Biomedical Ad Hoc grant (1345547) from the Burroughs Wellcome Fund. The content is solely the responsibility of the authors and does not necessarily represent the official views of the National Institutes of Health, any funding sources, or any affiliated institutions.

## Author Contributions

S.R.L., V.A.T., O.V.G., and R.W.F. together conceived the study.

S.R.L., M.P.R.S., and K.H. founded and co-directed CL-GSEC 2021 spring semester pilot under the supervision of O.V.G. and R.W.F.

D.M.C., V.A.T., and S.R.L co-directed the program in 2023-2024, including the acquisition of 2023 cohort program and evaluation data, under the supervision of O.V.G. and R.W.F.

S.R.L., V.A.T., D.M.C., R.A.G., O.V.G., and R.W.F. performed data analysis and interpretation.

R.W.F. acquired funding resources to support this project.

S.R.L., V.A.T., O.V.G., and R.W.F. together designed the figures and wrote the paper with input from all authors.

