## Supplementary File 1 for "A strong start for sustained success: inclusivity through a national group mentorship program for first-year graduate students"

**Supplementary File 1.** Descriptions of each CL-GSEC team, their general function, and their size/management.

| CL-GSEC Team | General Function | Description | Size/Management |
|---|---|---|---|
| Executive | Leadership | In charge of all internal operational aspects of CL-GSEC, including supporting each team in coordinating, organizing, and executing their roles and the overall team mission; as well representing CL-GSEC with the rest of Cientifico Latino, *Inc* and external organizations. | 2 Co-Directors |
| Recruitment | Logistics/Preparation Support | Recruit, screen, and match mentors and mentees in spring/summer for CL-GSEC program in academic year | Lead + 2 team members |
| Mentor-Mentee Relations | Mentee Resources | Main contact and manager of CL-GSEC mentorship small groups; liaison for conflict resolution | Lead + 2 team members |
| GradSchool 101 Workshops | Mentee Resources | Plan and host a series of panels covering general topics relevant for Grad School's first year students. | Lead + 2 team members |
| Community Engagement | Mentee Resources | Create a sense of community in the CL-GSEC cohort. Plan, execute, and host virtual social events (3x/year). Develop new community engagement initiatives (e.g., in person meet-ups). | Lead + 2 team members |
| Marketing | Operations Support | Advertise CL-GSEC events & announcements (including generating ads & social media) | Lead + 2 team members |
| Data | Operations Support | Maintain, organize, and analyze all the data gathered from CL-GSEC's operations. (application, pre-, post-, and check-in surveys) | Lead + 2 team members |
