## Supplementary File 2 for "A strong start for sustained success: inclusivity through a national group mentorship program for first-year graduate students"

**Supplementary File 2. Training resources for CL-GSEC scholars and mentors.**
The following is a document provided to CL-GSEC mentors and scholars at the start of the program.

### Scholar Resource: How to get the most out of your mentorship relationships

Oftentimes graduate students enter graduate school and place this program at the bottom of their commitments. Part of this program is that we expect scholars to honor the commitment being asked of them and find ways to prioritize these mentor-mentee group meetings. Below are ways to make sure that you are honoring the time commitment and honoring the commitment between yourself and your mentors.

#### Flexibility

Be flexible when scheduling your meeting. Learning to squeeze meetings back to back or in between classes is incredibly important.

- When filling out a when2meet make sure to fill it out with any possible free time you may have. Not just when you would prefer to have it. I understand it's inconvenient but this is going to happen a lot during graduate school
- Google Calendar will be your best friend for scheduling and keeping track of all your meetings while in grad school

#### Be Considerate

All of our mentors are also PhD students and are all volunteers. Their time is just as valuable as yours and it is extremely important to be considerate of their time.

- Be sure to respond to messages within a timely manner usually within 48 hours (see guidelines above)
- Do not ghost your mentor or skip meetings without an appropriate cancellation time. Not only does this impact the rest of your group but it is incredibly rude to do so. Things happen, but please try to notify your mentors ASAP if your schedule changes.

Effective communication between you and your mentors will be critical to a good mentor-mentee relationship. It's important to know that your mentor is not a mind reader and you cannot expect them to know what you need in order to succeed. Below are suggestions and examples of ways you can work on your own communication and interpersonal skills in order to help drive your relationship.

<u>Self Advocate</u>

One of the most important pieces of advice nobody tells you about grad school is that you must advocate for yourself. You have every right to ask for things that you believe you need and to speak up when you are struggling and need help. This is possibly the hardest lesson to learn and won't be easy.

<u>Be active and engaged during meetings</u>

- Give them your undivided attention and listen to both of your mentors during your meetings. Try not to take your meetings while you are in the lab or running an experiment (I know we're all guilty of this).
- Ask questions and offer your own feedback. These meetings should be more like a conversation and less of your mentors presenting to you; furthermore, giving feedback to mentors will go a long way in increasing the quality of mentorship.

<u>Aligning mentor and mentee expectations</u>

- Establish and communicate mutual expectations early in the relationship. Your mentor will go over their expectations, and you can and are encouraged to voice your own expectations.
- You can set these expectations over zoom or over an email. If you do not want to share your expectations with the group, you can email your mentors privately and voice your expectations or desires.
- Offer honest and open feedback on how the relationship is progressing and in turn, be open to feedback from your mentors.

<u>Create an agenda for meetings</u>

- We provide suggested topics for you and your mentors to follow for your monthly meetings. You can suggest other topics by providing your mentors with a tentative agenda 24 hours before your meeting.
    - You should create this agenda with your fellow mentees (this is also a great way to communicate with your fellow mentees outside of scheduled meetings and strengthen sense of community).

    - Don't be afraid to suggest specific topics, even if they may be somewhat niche! Worse case scenario, we encourage and expect your mentors to communicate with their fellow mentors in case a mentee needs advice related to an issue that the mentor may not be well versed in.

### Mentor Resource: Attributes for effective mentoring

Interpersonal skills and Communication

Effective communication between you and your mentees is critical to a good mentor-mentee relationship. It is important to know that your mentee may not know how to change their communication style or how to voice what they need. Below are examples of ways you can work on your own communication and interpersonal skills in order to help your mentee succeed.

- Actively listening to your mentee
    - Give them your undivided attention and listen to both words and the emotion behind the words
    - You don't have to offer advice. Often, just listening and acknowledging their struggles is effective.
- Aligning mentor and mentee expectations
    - Establish and communicate mutual expectations for the mentoring relationship. See "Communicating Expectations" section below for ways you can discuss this.
    - You should set expectations early on, and it may be helpful for you to get these in writing from your mentees.
        - Reach out to mentees individually to establish this if you have mentees with differing needs while establishing common expectations for the entire group as well
    - Offer honest and open feedback on how the relationship is progressing. Sometimes mentor-mentee relationships don't work and that is neither the mentor's fault or the mentee's fault. In cases like this, contact the Mentor-mentee team asap.
- **Don't be afraid to reach out to others**
    - Sometimes you don't have the answers to a question and that's ok. Be willing to reach out to other mentors and ask for help if a mentee has a situation or question you feel you can't answer.

Sponsorship and Career

Mentorship is not just about helping someone with the technical aspects of research. Sometimes it's about being there for support and motivation

- Help your mentee to celebrate the successes and offer support after failures.
    - One thing I often do with my mentees is see when they have finals and send words of encouragement during that time. I do this by adding it to my google calendar or you can even take a minute to schedule the message to send via slack.
    - For failures, I often listen and make sure they know that it is okay to fail. I often tell mentees about the time I failed an exam or botched an entire experiment. I'll let them know that science is all failure and offer ways that I deal with it.
- Developing mentee career self-efficacy
    - Encourage and foster your mentee's career aspirations. Don't judge them for it but try and help. I often times will send my mentees opportunities I think they may be interested in or encourage them to apply to things.
- Developing science identity
    - Academia can be very disruptive to one's identity in science.
    - Typically we say that we are graduate students and forget that we're in fact, scientists.
    - It's really important here to discuss ways your mentees wants to have their identity in science.
        - I often refer to myself as a woman in science or a first-generation student.
- Developing a sense of belonging
    - One of the largest parts of this program is to help encourage a sense of community and belonging.
- Promoting professional development
    - Identify opportunities for mentee professional development, and support their engagement in them
    - One way to do this is to just send them opportunities you see across twitter, your institution, word of mouth etc.
- Establishing and fostering mentee professional networks
    - If you're in the same field, try and introduce them to people you know either virtually or maybe at a conference.
- Actively advocating
    - This ties in above, vouch for them, and promote their work if possible. This can be done by sharing via Twitter or sharing their work with people who might be interested. Often times when I present, I make sure to promote the work my mentees did to contribute and promote their own projects that tie in.

Communicating Expectations

Here are some ideas for uncovering and clarifying your and your mentee's expectations:

- Share what you each expect from the relationship.
- Discuss the roles and responsibilities of each party.
- List any special needs or features that should be considered.
- Your mentee may not be comfortable discussing this in a group setting, so set up options for mentees to discuss or bring up suggestions privately (anonymous poll?)

Ask each other some critical questions:

- How much time, effort, and enthusiasm can you devote to this relationship?
- What do you think a mentor/mentee should do?
- Who's responsible for this relationship? What does that mean?
- Besides this relationship, what are your priorities?

Independently respond to the following and use your answers to start a conversation:

- What I expect to devote to this relationship is…
- I can give ____ time to this relationship.
- I anticipate meeting _____times a month.
- What I expect in terms of confidentiality / punctuality / communication is

### Ways to engage and connect between mentees and mentors:

Everything below is a list of suggestions for mentoring remotely.

- Communication
  - This can be done via email, slack, texts, whatsapp, etc etc. Figure out the best way to interact by asking your mentee
  - Do random check ins via your preferred communication. A weekly "Hey how's it going everybody"
    - You can do more structured check-ins like what's everyone's highlight of the week.
  - Send memes or videos!
  - Play video games!
  - Discord!
  - Things don't always have to be so structured between you.
- Monthly video meetings
  - These can be structured based on what is going on in the CL-GSEC program or based on what your mentees want to discuss
  - Create an agenda before meeting and share with your mentees so they can add things they want to discuss or prepare ahead of time
    - Create a Google poll for mentees

    - Start off each meeting by having everyone state a challenge or state a fun thing they did
    - After each meeting, you can discuss things your mentees want to discuss
- Have more relaxed social meetings
    - Coffee/tea hour
    - Game night check-in
