## Supplementary File 3 for "A strong start for sustained success: inclusivity through a national group mentorship program for first-year graduate students"

**Supplementary File 3. Suggested topics for monthly meeting discussions during a typical U.S. academic year.**

### September – Introduction, choosing rotations

i. What are some important things to assess during a rotation?
ii. What do you do if a rotation isn't going how you expected?
iii. What are some important things you need to ask a PI before you rotate?

### October – Balancing coursework, research and personal life/hobbies and mental health

**i.** On a scale from 1-10, how manageable do you think your coursework and lab work are?
**ii.** What percentage of your work time do you assign to coursework and lab work? What percentage do you assign to personal hobbies/life? Why?
**iii.** Do you feel you have enough time to pursue your own hobbies and other things that help you maintain good mental health?
**iv.** Is there anything we could do to help you manage to balance your work and lab work better?

### November – Finding and building community in graduate school

**i.** How do you identify and find mentors (other than your PI) in graduate school?
**ii.** What things are you passionate about outside of science?
**iii.** How do you get involved in groups outside of your lab in graduate school?

### December – First-semester check-in, hopes for second semester

**i.** What is something you are proud of in your first semester of graduate school?
**ii.** What is something you hope to do in your second semester?
**iii.** How to talk to PI's about time off during the holidays/in general?
**iv.** Were there any issues or struggles that you experienced that you want to go over or address (can reach out individually to mentors if not comfortable sharing with group)

### January – Choosing PI's/lab, asking PIs about funding

**i.** Do you see yourself thriving in the lab for the next 4-5 years?
**ii.** Other than your degree, what else are you looking to get out of your program? Will your lab of choice facilitate them?
**iii.** What are some of the best ways to ask about your future funding situation?

### February – Saying no to PI's, how to maintain good relationships with these PI's

i. Asking PIs to stay on as committee members, informal advisors
ii. How to talk to PIs that aren't your own (i.e. what to focus on when meeting with them, how often, etc.)

### March – Choosing thesis project, getting started in your lab

i. Identifying expectations between you and mentor in terms of getting started, freedom to pursue own project from scratch vs. building on previous one
ii. How to battle imposter syndrome as you start in the lab
iii. Collaborating/interacting with other students, post-docs etc. in the lab and asking for help/feedback

### April – Planning for fellowships, presenting at conferences/networking

**i.** Go over different types of available fellowships, funding agencies, what they entail etc.

ii. Use social media as a tool for networking, reaching out to people ahead of time or at conferences

#### May – Second-semester check-in, introduction to qualifying exam prep

i. Takeaways from the first year. What worked and what didn't?
ii. Encourage questions about the second year, getting involved in CL-GSEC, etc.
