## Supplementary File 4 for "A strong start for sustained success: inclusivity through a national group mentorship program for first-year graduate students"

**Supplementary File 4. Additional testimonials from CL-GSEC scholars and mentors.**

**SCHOLARS:**

*<u>What was the most positive part of your experience in the CL-GSEC program? What part did you find the most helpful?</u>*

“[For me, the] most positive part was getting together with the other mentees and mentors and [to] chat every month. Those interactions were the core of the program and I feel like they validated my experience as a grad student. Other mentees usually had similar experiences and we could talk our way through them, and mentors would often have useful feedback.”

“Having a smaller community within the Cientifico Latino community. I really enjoyed having the meetings because I could get advice from current graduate students.”

“My mentors were awesome and so were the other CL-GSEC mentees. After a few months it felt like I was catching up with old friends.”

“Seeing that I was not alone in struggling!”

“I appreciated hearing from other mentees and also from mentors about different grad school experiences. Also [I] found [it] helpful having someone outside of my bubble that I could reach out to.”

“Being able to openly talk about non-academic aspects of grad school. It’s been overwhelming to move across the country and start a grad program as a first Gen student.”

“The most positive part of my experience was having the opportunity to discuss the hardships and accomplishments of my first year of graduate school with a diverse group of scholars. As a Latina in STEM, it is often hard for me to find people who can understand and sympathize with my experiences so having a community of POC scholars allowed me to feel safe to express myself.”

“I found it helpful with the workshops that gave advice on important aspects of a first year grad student's experience - such as choosing the right mentor/PI. [It] was also good to talk to some graduate students who were familiar with funding opportunities for incoming PhD students like NSF GRFP and GEM.”

“The mentors!! They were so encouraging and it was nice to open up to people about mistakes/situations I was nervous about that I wasn't comfortable disclosing to people in my program.”

“Feeling like I wasn’t alone in this journey”

“I found a community I could identify with, that had experienced what [it] is like to come to the US to a PhD program as an international student. I received very good advice on how to choose a lab, apply to fellowships and navigate first year.”

“Knowing I wasn't alone in what I was going through and that I'm not the first or last person to experience hardships in grad school.”

“I think the most positive part was having a safe and supportive environment to share the struggles of my first year in a PhD program.”

“I found it helpful to listen to the workshops (whether recorded or live) because they brought up interesting questions that I did not consider or points that were very helpful to think about. I also appreciated that the workshops covered a wide range of topics and which were discussed with current students or alumni who have applicable experiences to share.”

“Having people to chat about the good, the bad, and the unexpected bits of graduate school was great for my mental health. Struggles were normalized, and we had great chats in a safe space about tackling these challenges. I am delighted with participating in this program and have recommended it to others!”

*What was your experience like with your mentors?*

“It was great having two mentors with different backgrounds. They were really helpful and were able to provide resources and adapt each meeting to our needs. They also provided a lot of support, validated our feelings as first years and provided tips on how to deal with imposter syndrome, stress etc.”

*Overall, how was your experience in the CL-GSEC program?*

“Good, though it was hard to participate at times because my first year felt so busy!”

“My experience in the CL-GSEC program was great! I looked forward to talking with my mentors and fellow mentees every month and talking about the new things I had accomplished. I also think we had a lot of discussions about doing what's best for you in grad school whether that be forming a better relationship with your advisor or taking days off.”

“It made me feel less alone.”

**MENTORS:**

*What was the most positive part of your experience in the CL-GSEC program?*

“Watching the mentees start to support each other”

“Getting to interact with mentees and feeling like we built a community together!”
“Getting to see the mentees encourage each other and seeing them actually talk to each other and reach out to us for help, meaning they trusted us”

“Getting to know my mentees and co-mentor was the best part!”

“Ability to meet people via this community and grow my network.”

*Overall, how was your experience in the CL-GSEC program?*

“I enjoyed the mentorship experience, our team had lots of venting sessions and I think it benefited them to see various PhD experiences.”

“I enjoyed being paired with a partner to help the burden of scheduling, or leading conversation. I really liked my mentees and that they all showed up for the group, and each other.”
